# Motivational learning biases are differentially modulated by genetic determinants of striatal and prefrontal dopamine function

**DOI:** 10.1101/2021.04.08.438916

**Authors:** Anni Richter, Lieke de Boer, Marc Guitart-Masip, Gusalija Behnisch, Constanze I. Seidenbecher, Björn H. Schott

## Abstract

Dopaminergic neurotransmission plays a pivotal role in appetitively motivated behavior in mammals, including humans. Notably, action and valence are not independent in motivated tasks, and it is particularly difficult for humans to learn the inhibition of an action to obtain a reward. We have previously observed that the carriers of the DRD2/ANKK1 TaqIA A1 allele, that has been associated with reduced striatal dopamine D2 receptor expression, showed a diminished learning performance when required to learn response inhibition to obtain rewards, a finding that was replicated in two independent cohorts. In the present study, we first report a replication of this finding in a third independent cohort of 99 participants. Interestingly, after combining all three cohorts (total N = 281), exploratory analyses regarding the COMT Val108/158Met polymorphism suggest that homozygotes for the Met allele, which has been linked to higher prefrontal dopaminergic tone, show a lower learning bias. Our results corroborate the importance of genetic variability of the dopaminergic system in individual learning differences of action-valence interaction and, furthermore, suggest that motivational learning biases are differentially modulated by genetic determinants of striatal and prefrontal dopamine function.

## Introduction

The impact of motivation on cognitive functions has been subject to intense investigation over the past two decades. While the influence of motivational salience on cognitive processes and goal-directed behavior is common knowledge nowadays, theories of instrumental learning have until recently neglected the influence of outcome valence on action initiation. However, when action and valence are experimentally orthogonalized, signals that predict reward are prepotently associated with behavioral activation, whereas signals that predict punishment are intrinsically coupled to behavioral inhibition. This finding has been robustly replicated in multiple studies [1-15]. Understanding the neurocognitive mechanisms underlying this behavioral bias is thus important for developing more comprehensive theories of instrumental learning.

Numerous studies in a multitude of species, including humans, indicate the importance of dopamine (DA) in the neural manifestation of motivated behavior and the human dopaminergic system is subject to considerable genetic variability. According to a prevalent view in reinforcement learning and decision making, DA neurons signal reward prediction errors [16-18], in the form of phasic bursts for positive prediction errors and dips below baseline firing rate for negative prediction errors [19], resulting in corresponding peaks and dips of DA availability in target structures, most prominently the striatum [20-23]. In the striatum, increased DA release in response to an unexpected reward reinforces the direct pathway via activation of D1 receptors and thereby facilitates the future generation of *go* choices under similar circumstances, while dips in DA levels in response to an unexpected punishment reinforce the indirect pathway via reduced activation of D2 receptors, thereby facilitating the subsequent generation of *no-go* choices in comparable situations [24-27].

In line with those assumptions, we observed in a previous study [5] that the coupling of action and valence during learning was modulated by a genetic variant linked to striatal DA D2 receptor expression. We argued that A1 carriers with presumably less D2 receptors would be assumed to have less limitation of dopaminergic signaling after negative prediction errors in the indirect pathway and a shift to a more action-oriented behavioral pattern mediated by the direct pathway. In line with that framework, in a recent study, de Boer et al. [10] found a positive correlation between the strength of the action by valence interaction and dorsal striatal D1 receptor availability measured using positron emission tomography (PET). Therefore, striatal dopaminergic effects may be sufficient to explain biased motivational learning [9,10]. On the other hand, Guitart-Masip et al. [4] observed that levodopa administration led to a reduced coupling of action and valence that cannot be explained by striatal action of DA. The authors attributed their observation to an effect on prefrontal cortex (PFC) functioning, where DA plays a role in facilitating working memory and attentional processes [28-30] that may help to overcome the biased behavior. This effect of levodopa administration was recently replicated in patients with non-tremor Parkinson’s disease [14], and studies investigating frontal network dynamics using electroencephalography further demonstrate that prefrontal control processes (as indexed by higher mid-frontal theta power) are important to overcome biased behavior [1,8]. Therefore, DA may influence these learning biases in a regionally specific manner.

Numerous previous studies have investigated the influence of candidate single nucleotide polymorphisms (SNPs) of DA on instrumental learning [25,31-34]. As the expression of several key molecules of the dopaminergic system shows a characteristic regional distribution in the brain, genetically mediated differences may also provide some information about the contributions of different brain regions to DA-dependent learning and memory processes [34-36]. In the current study, we aimed to examine differential contributions of two prominent dopaminergic SNPs: the DRD2/ANKK1 TaqIA SNP (rs1800497) that has been implicated in striatal DA metabolism and the COMT Val108/158Met SNP (rs4680) which has been shown to influence prefrontal DA availability.

The TaqIA polymorphism has repeatedly been linked to lower striatal D2 binding availability using PET in carriers of the less common A1 allele [37-40]. With respect to motivated behavior, Stice et al. [41] found stronger midbrain activation in A1 carriers compared with A2 homozygotes on reward expectancy, and Stelzel et al. [42] reported generally increased striatal BOLD signaling in A1 carriers. In addition, relative to A2 homozygotes, A1 carriers showed poorer performance in avoiding actions associated with punishment and lower activations of PFC and striatum during processing of negative feedback [31-33]. Catechol-O-methyltransferase (COMT) plays a key role in the breakdown of DA in the PFC [43,44]. The frequent Val108/158Met SNP in the *COMT* gene (chromosome 22) leads to an amino acid exchange from valine (Val) to methionine (Met). In Met carriers reduced enzymatic activity and increased prefrontal DA availability have been observed, presumably due to lower thermostability of the enzyme [45].. This SNP has mainly been investigated with respect to PFC-dependent executive functions (for reviews, see [46,47]), and a meta-analysis of functional magnetic resonance imaging (fMRI) studies confirmed that Met-carriers show more efficient performance in executive functions and higher neural activations during emotion processing [36]. In the context of motivated behavior, the Met allele has been associated with more successful reward learning (for a meta-analysis see [34]). Moreover, Met allele carriers adapt behavior more rapidly on a trial-to-trial basis during reinforcement learning [25,31].

We have previously shown in two independent cohorts that carriers of the A1 allele of the DRD2/ANKK1 TaqIA polymorphism show a rather selective deficit in learning to inhibit an action to receive a reward [5]. With our present study we followed two aims: Firstly, we aimed to replicate our finding on the TaqIA polymorphism in a third independent cohort and to investigate the nature of the genetic effects more closely using trial-by-trial behavioral analysis and computational modeling in the combined dataset (N=281). Secondly, we aimed to assess a potentially modulatory role of prefrontal DA availability, using the widely studied COMT Val108/158Met polymorphism as a proxy. Regarding the TaqIA SNP, we hypothesized that, in line with our previous observations [5], A1 carriers would show a higher coupling of action and valence. With respect to the COMT polymorphism, we hypothesized that, given the preferential role of COMT in PFC versus striatal DA availability, carriers of the low-activity Met allele would more readily overcome the learning bias and show less coupling of valence with action.

## Materials and methods

### Participants

In addition to our previously described two cohorts of 87 and 95 participants [5], 99 newly recruited participants were tested (55 females and 44 males; age: range 20–34 years, mean 25.2 years, SD = 2.6 years; demopgraphic description of all three samples in Supplementary Table S1). According to self-report all participants were of European ethnicity, right-handed, had obtained at least a university entrance diploma (Abitur) as educational certificate, had no present or past neurological or mental disorder, alcohol or drug abuse, did not use centrally acting medication, and had no history of psychosis or bipolar disorder in a first-degree relative. Additionally, given the design of the experiment, regularly gambling was defined as an exclusion criterion for participation.

All participants gave written informed consent in accordance with the Declaration of Helsinki and received financial compensation for participation. The study was approved by the Ethics Committee of the Faculty of Medicine at the Otto von Guericke University of Magdeburg.

### Genotyping

Genomic DNA was extracted from blood leukocytes using the KingFisher™Duo Prime Purification System (Thermo Scientific(tm)) according to the manufacturer’s protocol. Genotyping of the SNPs DRD2/ANKK1 TaqIA (NCBI accession number: rs1800497) and COMT Val108/158Met (rs4680) was performed using PCR-based restriction fragment length analysis according to previously described protocols [5,35,48-50]. A1 carriers of the TaqIA SNP were grouped together (A1+: A1/A1 and A1/A2; A1-: A2/A2) as in previous studies [5,31-33,41,42,48,49].

### Paradigm

We used a previously employed *go/no-go* learning task with orthogonalized action requirements and outcome valence [3]. Detailed descriptions of the task have been presented previously [5,6]. Figure 1A displays the trial timeline. Briefly, each trial consisted of the presentation of a fractal cue, a target detection task, and a probabilistic outcome. First, one out of four abstract fractal cues was displayed. Prior to the beginning of the task, participants were informed that a fractal indicated i) whether they would subsequently be required to perform a target detection task by pressing a button (*go*) or not (*no-go*) and ii) the possible valence of the outcome of the subjects’ behavior (reward/no reward or punishment/no punishment). Importantly, subjects were not instructed with respect to the contingencies of each fractal image and had to learn them by trial and error. There were four trial types: press the correct button in the target detection task to gain a reward of 0.50 € [“*go to win*” (*gw*)]; press the correct button to avoid a punishment of -0.50 € [“*go to avoid losing*” (*gal*)]; do not press a button to gain a reward [“*no-go to win*” (*ngw*)]; do not press a button to avoid punishment [“*no-go to avoid losing*” (*ngal*)]. The outcome was probabilistic (see figure 1B). To avoid incidental effects of specific cue images, the association of the fractal images with the specific conditions (go vs. no-go * reward vs. punishment) was randomized across participants. The task included 240 trials (60 trials per condition) and was divided into four sessions. Subjects were told that they would be paid their earnings of the task up to a total of 25 € and a minimum of 7 €. Before starting the actual learning task, subjects performed 10 trials of the target detection task in order to familiarize themselves with the speed requirements.

**Fig. 1.**
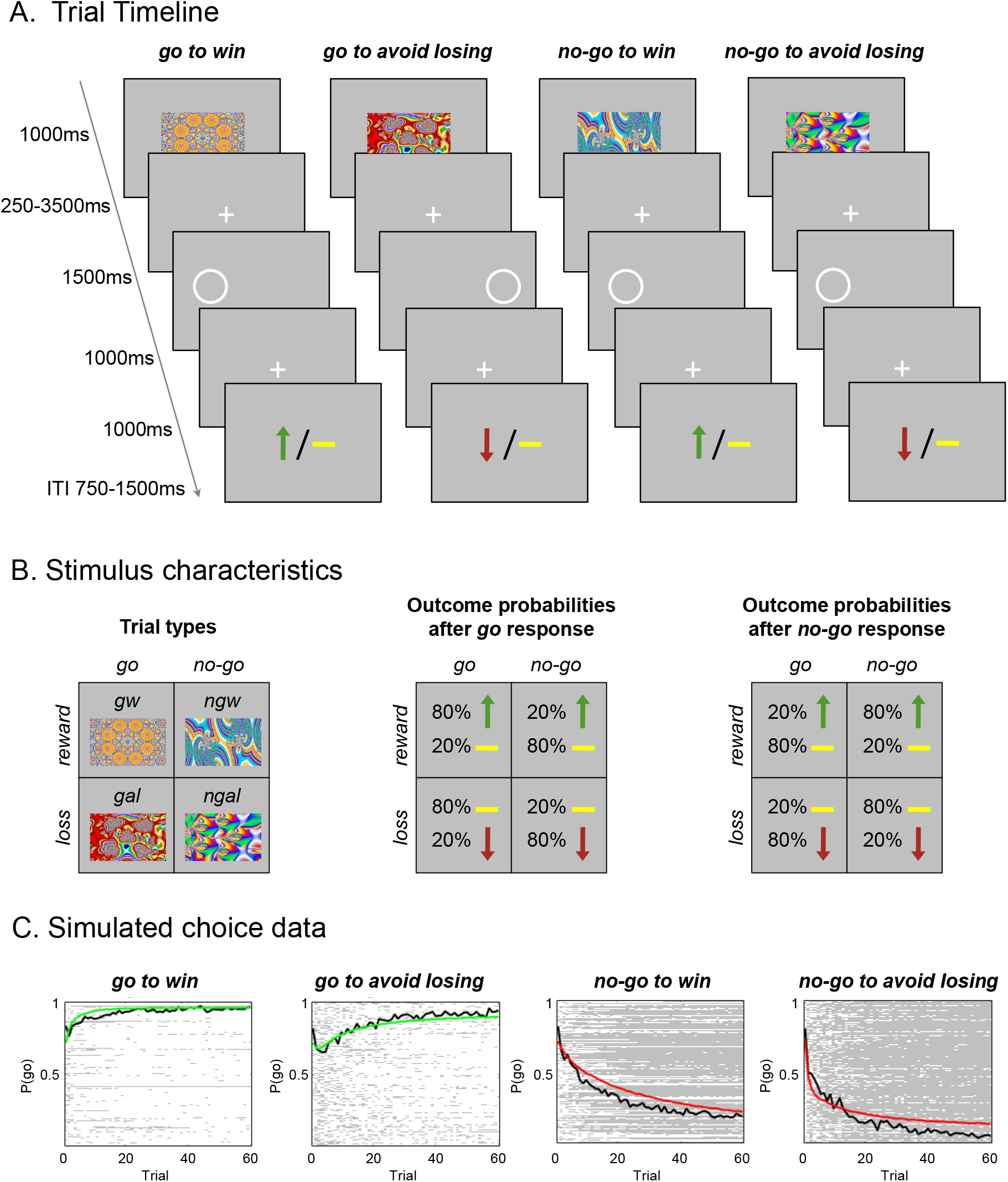
Experimental paradigm and participant performance. *(A)* Probabilistic monetary *go/no-go* task. Fractal cues indicate the condition - a combination of action (*go* or *no-go*) and valence (*reward* or *punishment*). On *go* trials, subjects press a button for the side of a circle. On *no-go* trials they withhold a response. Arrows indicate *rewards* (green) or *punishments* (red). Horizontal bars (yellow) symbolize the absence of a *reward* or *punishment*. ITI, intertrial interval. *(B)* The schematics represent for each condition the nomenclature (left), the possible outcomes and their probabilities after a *go* response (middle), and the possible outcomes and their probability after a *no-go* response (right). *gw*: *go to win*, gal: *go to avoid losing*, ngw: *no-go to win*, ngal: *no-go to avoid losing. (C)* Simulated choice data according to the model parameters of the winning model. Colored lines represent the simulated group mean probability of performing a *go* on each trial (green for *go* conditions, where *go* is the correct response; red for *no-go* conditions, where *no-go* is the correct response). Black lines indicate the group mean for participants’ actual *go* responses on each trial. In the plot area, each row represents one participant’s choice behavior for each trial (281 x 60 pixels). A white pixel reflects that a participant chose *go* on that trial; a gray pixel represents *no-go*. Participants made more *go* responses to *win* vs. *avoid losing* cues, reflecting the motivational bias. Overall, they successfully learned whether to make a *go* response or not (proportion of *go* responses increases for *go* cues and decreases for *no-go* cues). Figures *(A)* and *(B)* adapted from Richter et al. [5].

### Statistical analysis of accuracy

Accuracy was analyzed using IBM^®^ SPSS^®^ Statistices version 21. The percentage of correct choices in the target detection task (button press in *go* trials and omission of responses in *no-go* trials) was collapsed across time bins of 30 trials per condition. To assess the learning enhancement, the slope was calculated by substracting the mean values in the first half of the experiment from the mean values of the second half of the experiment (slope = mean[2^nd^ half] - mean[1^st^ half]).

For the replication of our previous study [5] in the new cohort (N=99) we compared TaqIA genotype groups with a *t*-test for independent samples and investigated task effects with a mixed analysis of variance (ANOVA) with time (1^st^/2^nd^ half), action (*go/no-go*), and valence (*win*/*avoid losing*) as within-subject factors.

Then, by combining all three datasets (N=281), we included the two genotypes as between-subject factors in the analysis and added cohort (three cohorts represented in two dichotomous dummy coded variables for cohort 2 and 3), age and gender as covariates (analysis of covariance, ANCOVA). The increased number of participants allowed us to run a logistic regression on the trial-by-trial *go* responses as in Swart et al. [9] which more accurately analyzes the data, as it is closer to the actual behavior of each participant by including inter- and intraindividual variability (see supplementary methods for details).

Unless stated otherwise, independent samples *t*-tests were used as *post hoc* tests, and the significance threshold was set to .05, two-tailed. Whenever Levene’s test was significant, statistics were adjusted, but for better readability, uncorrected degrees of freedom are reported.

### Computational Modeling of task performance

Computational Modeling of task performance was employed using MATLAB^®^ R2016B (Mathworks^®^). We used a previously published modeling procedure [3,51]. Detailed descriptions of the reinforcement learning models as well as the model fitting procedure and comparison have been described in a recent study of age effects in the same task [6]. Briefly, we constructed six nested reinforcement learning models to fit participants’ behavior (Table 2). The base model was a Q-learning algorithm [52] that used a Rescorla-Wagner update rule to independently track the action value of each choice (*go; no go*), given each fractal image, with a learning rate (*ε*) as a free parameter. In this model, the probability of choosing one action on a trial was a sigmoid function of the difference between the action values scaled by a slope parameter that was parameterized as sensitivity to reward (*ρ)*. This basic model was augmented with an irreducible noise parameter (*ξ*) and then further expanded by adding a static bias parameter to the value of the *go* action (*b*). Further, we allowed for separate sensitivities to rewards (*ρ*_*win*_) and punishments (*ρ*_*lose*_). As in our recent study of age effects [6], the model was then extended by adding a constant Pavlovian value of 1 or -1 to the value of the *go* action as soon as the first reward for *win* cues or the first punishment for *avoid losing* cues, respectively, was encountered. This fixed Pavlovian value was weighted by a further free parameter (Pavlovian parameter) into the value of the *go* action (*π*). Model comparisons demonstrated a better fit compared to a variable Pavlovian value used in previous studies [1,3,10] (see Table 2). As in previous reports [3,51], we employed a hierarchical Type II Bayesian procedure using maximum likelihood to fit simple parameterized distributions for higher-level statistics of the parameters. All six computational models were fit to the data using a single distribution for all participants. This fitting procedure was, therefore, blind to the existence of different genotype groups with putatively different parameter values. Models were compared using the integrated Bayesian Information Criterion (iBIC) with small iBIC values indicating a model that fits the data better after penalizing for the number of data points associated with each parameter. Finally, we assessed genotype-related effects on all modeling parameters using IBM^®^ SPSS^®^ Statistices version 21. To test for differences regarding specific model parameters we calculated *t*-tests for independent samples. As one could not exclude that not one specific parameter but a combination of them differed between genotypes, we performed a multivariate test of differences – a linear discriminant analysis (LDA). The purpose of LDA was to find a linear combination of the six model parameters that gives the best possible separation between the genotype groups. This method simultaneously accounts for differences in combinations of variables between groups over and beyond differences across single multiple variables [53].

## Results

### Reduced learning performance in DRD2/ANKK1 TaqIA A1 carriers

In our previous study [5] we observed that in the *no-go to win* condition TaqIA A1 carriers showed a significantly diminished improvement from the first to the second half of the experiment compared to A2 homozygotes (cohort 1: *t*_85_ = -2.78, *p* = 0.007; cohort 2: *t*_93_ = -2.16, *p* = 0.033). As expected, we replicated this finding in our current sample (cohort 3: *t*_97_ = 2.05, *p* = .043; Figure 2A). In all other conditions A1 carriers and A2 homozygotes did not significantly differ (all *p* > .100), nor in gender (*p* = .621), age (*p* = .749), the number of smokers and nonsmokers (*p* = .084) or in the COMT Val108/158Met genotype distribution (*p* = .901).

**Fig. 2.**
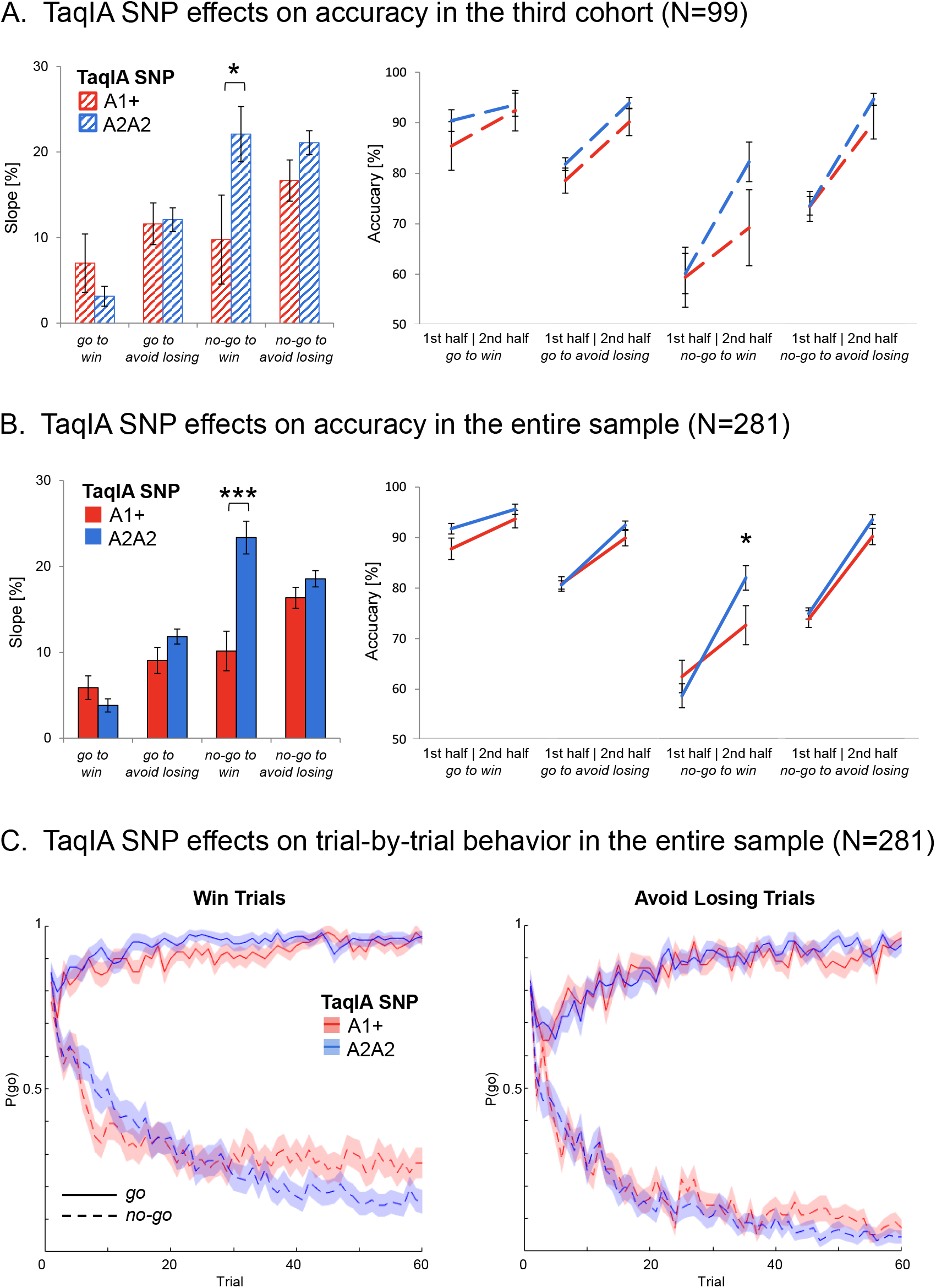
Effects of DRD2/ANKK1 TaqIA genotype on choice performance. *(A)* and *(B)* Effects of DRD2 TaqIA genotype on choice performance in the third cohort (N = 99) and in the entire sample (N = 281). Compared to the A2 homozygotes, A1 carriers showed a diminished learning to withhold an action to receive a reward. Left panels: Bar plots show mean differences between correct response rates (±SEM) during second half versus the first half of trials for each condition. This score represents the observed four-fold interaction of *action* x *valence* x *time* x *genotype*. Right panels: Line charts show mean values of correct responses (±SEM) in the first and the second half of trials for all four conditions. *Post hoc* comparisons via *t*-tests: \**p* < 0.05, \*\*\**p* < 0.001. *(C)* Trial-by-trial proportions of *go* responses (±SEM) to *go* cues (solid lines) and *no-go* cues (dashed lines) across cue types. *Win* and *avoid losing* condition seperately and colors depict TaqIA genotypes. TaqIA A1 carriers showed an enhanced effect of cue valence on *go* responding especially in the *no-go to win* condition with further progress of the experiment (lines are mostly sperated). Adapted scripts of Swart et al. [9] were used to generate figures.

Furthermore, we also analyzed task effects and replicated previous results showing an action by valence interaction on overall task performance [1-15] (see supplementary results and Table S2 for details).

### DRD2/ANKK1 TaqIA and COMT genotypes differentially modulate motivational learning biases

Our further analyses of genetically driven effects were performed in the entire sample comprising all three cohorts (N = 281 participants). Genotype frequencies were in Hardy–Weinberg equilibrium (all *p* > .145), and there was no linkage between the two polymorphisms (*p =* .971; for detailed demographics see Table 1).

**Table 1.** Descriptive data of the entire sample regarding TaqIA and COMT genotypes.

| DRD2 TaqIA |  | A1+ | A1- | A1+ > A1- |  |  |
| --- | --- | --- | --- | --- | --- | --- |
| Gender (N | | | | $\chi^2 = 3.46$ | | |
| Women/Men) | | 44/55 | 102/80 | $p = .063$ | | |
| Age in years | | | | $t_{279} = 1.30$ | | |
| (M +/- SD) | | 25.1 +/- 3.1 | 24.6 +/- 2.6 | $p = .195$ | | |
| Non-Smokers/ | | | | $\chi^2 = 2.16$ | | |
| Smokers (N) | | 70/29 | 143/39 | $p = .141$ | | |
| COMT (N | | | | $\chi^2 = 0.06$ | | |
| MM/VM/VV) | | 30/45/24 | 53/83/46 | $p = .971$ | | |
| COMT | MM | VM | VV | MM > VV | VM > VV | MM > VM |
| Gender (N | | | | $\chi^2 = 0.59, p = .743$ | | |
| Women/Men) | 43/40 | 64/64 | 39/31 |  |  |  |
| Age in years | | | | $t_{151} = 1.36$ | $t_{196} = 1.37$ | $t_{209} = 0.09$ |
| (M +/- SD) | 25.0 +/- 2.7 | 24.9 +/- 2.8 | 24.3 +/- 3.0 | $p = .176$ | $p = .174$ | $p = .931$ |
| Non-Smokers/ | | | | $\chi^2 = 1.90, p = .387$ | | |
| Smokers (N) | 63/20 | 93/35 | 57/13 |  |  |  |
| TaqIA (N | | | | $\chi^2 = 0.06, p = .971$ | | |
| A1+/A1-) | 30/53 | 45/83 | 24/46 |  |  |  |

Demographic data are pooled across all three cohorts (cohort 1 and 2 from Richter et al. [5], and the newly investigated cohort 3). N = number, M = mean, SD = standard deviation, VM: Val/Met heterozygotes, VV: Val homozygotes, A1+: carriers of the A1 allele, A1-: A2 homozygotes.

**Table 2.** Integrated Bayesian Information Criteria (iBIC) for tested models

| Model no. | Model parameters | No. of parameters | Likelihood | Pseudo-R<sup>2</sup> | iBIC |
| --- | --- | --- | --- | --- | --- |
| 1 | $\varepsilon, \rho$ | 2 | -23463 | 0.498 | 46970 |
| 2 | $\varepsilon, \rho, \xi$ | 3 | -23314 | 0.501 | 46695 |
| 3 | $\varepsilon, \rho, \xi, b$ | 4 | -21798 | 0.534 | 43685 |
| 4 | $\varepsilon, \rho_{win}, \rho_{lose}, \xi, b$ | 5 | -21334 | 0.544 | 42779 |
| 5 | $\varepsilon, \rho_{win}, \rho_{lose}, \xi, b, \pi_{variable}$ | 6 | -21137 | 0.548 | 42406 |
| **6** | **$\varepsilon, \rho_{win}, \rho_{lose}, \xi, b, \pi_{constant}$** | **6** | **-21106** | **0.549** | **42346** |

Boldface type: winning model statistics, $\varepsilon$ : learning rate, $\rho_{win}$ : weighting of reward on win trials, $\rho_{lose}$ : weighting of punishments on lose trials. $\xi$ : irreducible noise, $b$ : go bias, $\pi$ : Pavlovian bias, iBIC: integrated Bayesian information criterion (smaller iBIC values indicate a better model fit). Descriptives for the parameters in the winning model (M +/- SD): $\varepsilon = 0.26 \pm 0.15$ , $\rho_{win} = 15.32 \pm 13.30$ , $\rho_{lose} = 7.51$ $\pm 4.03$ , $\xi = 0.96 \pm 0.06$ , $b = 1.10 \pm 0.74$ , $\pi_{constant} = 0.65 \pm 0.57$ .

In line with our previous work [5], we observed for the TaqIA SNP a significant *genotype* x *time* x *action* x *valence* interaction (*F*_1,271_ = 11.18, *p* = .001; see Figure 2B), as well as significant interactions of *genotype* x *time* (*F*_1,271_ = 11.08, *p* =.001) and *genotype* x *time* x *action* (*F*_1,271_ = 11.94, *p* = .001). *Post-hoc* comparisons revealed that A1 carriers exhibited an overall significantly worse learning performance throughout the experiment compared to A2 homozygotes (overall slope: *t*_279_ = -3.72, *p* < .001, Cohen’s *d* = 0.47). This effect was solely carried by the *no-go* conditions (*no-go* slope: *t*_279_ = -4.56, *p* < .001, Cohen’s *d* = 0.58; *go* slope: *p* = .748), and specifically by the *no-go to win* condition (*ngw* slope: *t*_*279*_ = - 4.41, *p <* .001, Cohen’s *d* = 0.54; all other conditions: all *p* > .087). As displayed in Figure 2B and 2C, the TaqIA A1 carriers reached their learning asymptote earlier and to a lower level. They significantly differed in performance from the A2 homozygotes only during the second half of the experiment, pointing to different learning capacities (overall 2^nd^ half: *t*_279_ = -2.21, *p* = .028, Cohen’s *d* = 0.35; *no-go* 2^nd^ half: *t*_279_ = -2.28, *p* = .024, Cohen’s *d* = 0.29; *ngw* 2^nd^ half: *t*_279_ = -2.06, *p* = .041, Cohen’s *d* = 0.26; equivalent 1^st^ half comparisons: all *p* > .340). A summary of the statistics is displayed in Supplementary Tables S3 and S4.

The combined datasets allowed for a logistic regression on the trial-by-trial *go* responses. This analysis confirmed the ANCOVA results with A1 carriers showing significantly diminished *no-go to win* performance in the course of the experiment (see Figure 2C and supplementary results for details).

For the COMT Val108/158Met polymorphism, we observed a trend towards a significant four-way interaction *genotype* x *time* x *action* x *valence* (*F*_2,271_ = 2.96, *p* = .053). Met homozygotes showed significantly increased learning throughout the experiment in the *no-go to win* (*ngw* slope: *t*_209_ = 2.02, *p* = .045; Figure 3) and the *go to avoid losing* conditions (*gl* slope: *t*_209_ = 2.48, *p* = .014) compared to heterozygotes (other conditions: all *p* > .922). The logistic regression did not show an effect of COMT genotype (*p* = .381; see supplementary results and Figure S2 for details).

**Fig. 3.**
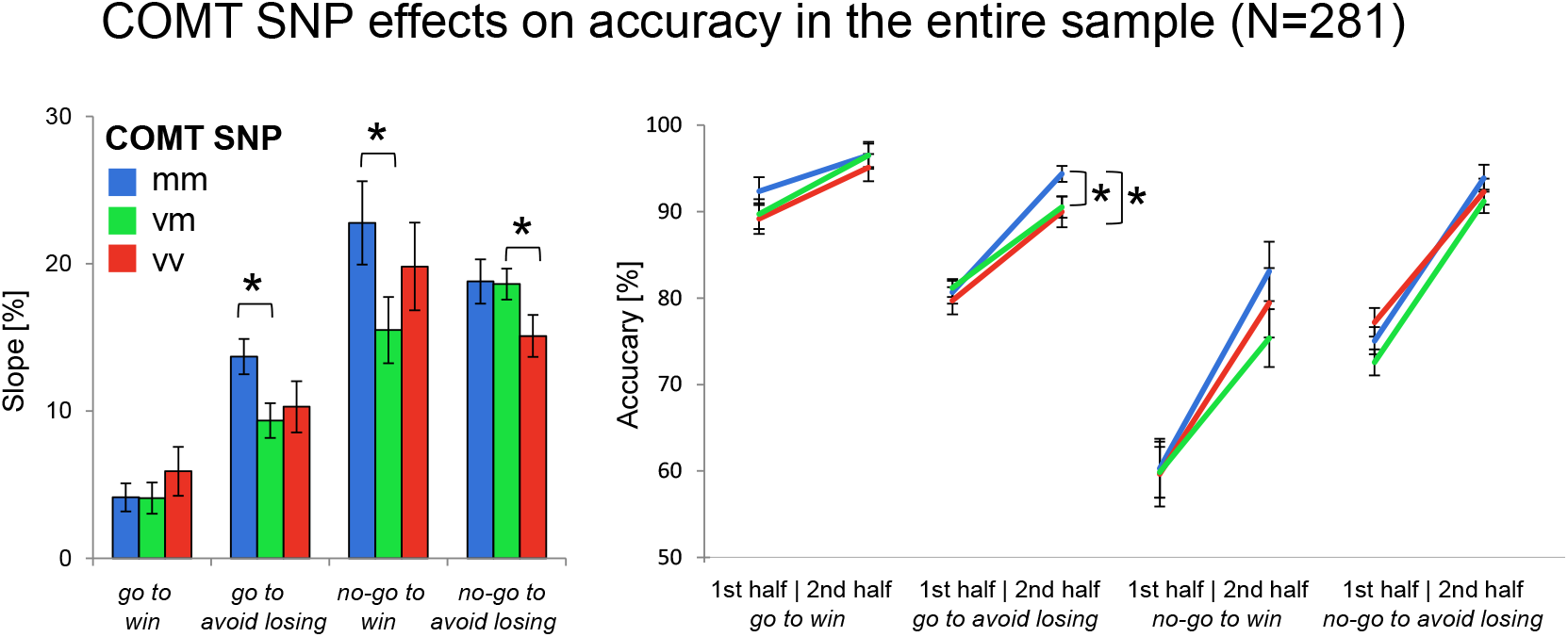
Effects of COMT genotype on choice performance in the entire sample. Left panels: Bar plots show mean differences between correct response rates (±SEM) during second half versus the first half of trials for each condition. This score represents the observed four-fold interaction of *action* x *valence* x *time* x *genotype*. Right panels: Line charts show mean values of correct responses (±SEM) in the first and the second half of trials for all four conditions. Met homozygotes showed increased learning throughout the experiment in the *no-go to win* and *go avoid losing* condition relative to heterozygotes. *Post hoc* comparisons via *t*-tests: \**p* < 0.05.

In light of previous evidence that Met homozygotes have a higher response bias relative to Val carriers [34,54-56], in an additional analysis participants were separated into Met homozygotes (Met/Met) and Val allele carriers (Val/Val and Val/Met). The ANCOVA revealed a significant *genotype* x *time* x *action* x *valence* interaction (*F*_1,273_ = 4.30, *p* = .039) as well as a significant main effect of COMT genotype (*F*_1,273_ = 4.55, *p* = .034) and interestingly also a significant interaction of the COMT with the TaqIA genotype (*F*_1,273_ = 3.88, *p* = .050). The latter finding indicates a benefical effect of Met homozygosity on overall performance in A1 carriers (*t*_97_ = 2.31, *p* = .024) but not in A2 homozygotes (*p* = .971).

We controlled for potential effects in reaction times (participants were explicitly instructed to respond accurately) and false responses in the target detection task (i.e., left when the target was on the right side of the display or vice versa) and found no significant differences between genotype groups (*p* > .187; see supplement for details).

### Computational Modeling of Task Performance

To identify components of the observed asymmetry during learning, we constructed six nested reinforcement learning models to fit participants’ behavior (Table 2). Our computational modeling approach demonstrated that the marked asymmetry in learning could be best accounted for by the model including separate parameters for sensitivity to rewards and punishments as well as a learning rate, an irreducible noise parameter, a constant *go* bias parameter, and a constant Pavlovian bias parameter (see Table 2), which is consistent with our recently published lifetime study on motivational learning [6]. The simulations of the winning model are presented in Figure 1C. Neither one specific model parameter (independent samples *t*-tests: all *p* > .119), nor a linear combination of the parameters (LDA: all *p* > .636) showed significant genotype-related differences.

## Discussion

In the present study, we investigated how genetic determinants of striatal and prefrontal DA function modulate learning biases when action and valence are experimentally orthogonalized. Using the previously established valenced *go/no-go* task [3], we provide independent confirmation for a selective deficit of DRD2 TaqIA A1 carriers in learning to inhibit an action in order to obtain a reward. Moreover, our exploratory analysis yielded preliminary evidence that COMT Met homozygotes show superior learning during trials with incongruent coupling of action and valence.

### Genetically driven contributions to the coupling of action and valence during learning

For the TaqIA polymorphism, we replicated our previous observation [5] that A1 carriers show a stronger coupling of action and valence in a third independent cohort. As in our previous study, A1 carriers exhibited a specific impairment in learning to withhold actions in reward contexts. When combining all three datasets (N = 281), we could more closely investigate the nature of this effect. Moreover, the larger sample size of our three combined samples made it possible to investigate the effects of and potential interactions with the COMT Val108/158Met polymorphism.

Due to previous knowledge about their neurophysiological consequences, the genetic polymorphisms studied here allow conclusions about differential contributions of striatal and prefrontal DA function to instrumental control mechanisms [34-36]. D2-type DA receptors are primarily expressed in the striatum (*post mortem* autoradiography: [57-59]; *in vivo* PET: [60,61]). They function as both postsynaptic inhibitory receptors and as presynaptic autoreceptors that regulate neurotransmission via negative feedback ([62], for reviews, see [63,64]). While DRD2 is, albeit sparsely, expressed in extrastriatal regions (2-8% of the expression level in the striatum [65]) and cortically mediated effects can thus not be excluded, differences for the ANKK1 TaqIA genotypes have thus far only been observed for the striatum - with lower DRD2 expression resp. binding availability in A1 carriers (*post mortem* autoradiography: [66-68]; *in vivo* PET: [37-40]).

In contrast, the decreased enzymatic activity of COMT in 108/158Met homozygotes primarily affects DA availability in the PFC, which has been attributed to the sparse cortical expression of the DA transporter (DAT) [45,69]. Therefore, the COMT polymorphism has mostly been studied in relation to PFC-dependent executive functions (for reviews, see [46,47]; for a meta-analysis see [36]). With respect to motivated behavior, homozygosity for the Met allele has been associated with relatively increased reward learning (for a meta-analysis see [34]). In our study, Met homozygosity was associated with stronger learning enhancement during Pavlovian conflict (i.e., incongruent coupling of action and valence) throughout the experiment. Our data suggest that higher prefrontal DA levels may improve performance when motivational biases are involved. Guitart-Masip et al. [4] hypothesized this mechanism to explain their unexpected finding that levodopa administration led to a reduced coupling of action and valence. These surprising effects of levodopa were replicated in a recent study [14]. Moreover, electrophysiological studies [1,8] point to the involvement of the same prefrontal control mechanism, when subjects learn to overcome Pavlovian conflicts.

Although COMT activity is of negligible importance to striatal DA availability [70], a potential indirect effect of COMT on striatal DA function cannot be excluded. Animal studies suggest that transgenic mice with increased COMT activity, equivalent to the relative increase in activity observed with the human COMT Val allele, do not only show deficits in PFC-dependent tasks (e.g., stimulus–response learning and working memory), but also increased DA release capacity in the striatum [71]. This finding corroborates earlier human neuroimaging studies that reported higher midbrain DA synthesis capacity in Val compared to Met homozygotes [72,73]. Thus, the effects of the COMT polymorphism on motivational learning may not only be explained by increased DA signaling in the cortex, but also contain a minor component of presynaptic DA availability in the striatum.

## Limitations

A limitation in the interpretation of our data that is also common in other studies on this topic lies in the fact that the molecular mechanisms underlying the observed effects are still under debate. It is well known that the TaqIA polymorphism is not located within the DRD2 gene but 10kb downstream of its termination codon on chromosome 11q23.1, within the coding region of the adjacent ankyrin repeat and kinase domain containing 1 (*ANKK1*) gene [74,75]. The molecular mechanisms underlying the effects of ANKK1 TaqIA on striatal DRD2 availability have not been conclusively established. Multiple mechanisms have been proposed, including linkage disequilibrium [49,67,76-78] or a potential direct interaction of ANKK1 with the D2 receptor at protein level, potentially modulated by the TaqIA polymorphism [79-81] (for a review, see [82]; see Supplementary Discussion for details). Similarly, for the COMT Val108/158Met polymorphism, it remains to be determined how COMT-dependent DA inactivation in brain regions with low DAT expression is realized. There is only limited evidence for extracellular activity of membrane-bound COMT [83], and the predominant evidence points to intracellular orientation and activity, requiring a DAT-independent uptake mechanism [44,84] (see Supplementary Discussion).

A further limitation lies in our modeling approach, which failed to reflect the very robust and replictad effect of the DRD2 TaqIA SNP on learning gain throughout the experiment in the *no-go to win* condition and on the time-dependent valence effect on individual *go/no-go* responses. One explanation could be that the model space does not include the computational mechanism to differentiate, for example, instrumental from Pavlovian contributions. This should be addressed in future studies.

## Conclusion

It is not clear how differential effects of striatal and prefrontal DA function contribute to motivational learning biases. With our study, we demonstrate by assesing the contributions of two well-studied genetic polymorphisms that DRD2/ANKK1 TaqIA A1 carriers with presumably fewer striatal D2 receptors and less limitation of striatal dopaminergic signaling after negative prediction errors in the indirect pathway showed a shift to a more action-oriented and biased behavioral pattern. COMT Val108/158Met Met homozygotes, who presumably exhibit higher prefrontal DA activity, showed less biased learning, possibly reflecting more efficient frontal control.

## Supporting information

Supplemental Methods, Results and Discussions

## Acknowledgements

We are grateful to Herta Flor for valuable comments on the manuscript. We thank Iris Mann, Catherine Libeau and Timo Lemme for help with testing.

## Author contributions

AR, LdB, MG-M, CIS, and BHS wrote the manuscript. AR, MG-M and BHS conceptualized the study design. AR and GB collected the data. AR, LdB and GB analyzed and curated data.

