## Supplemental Methods, Results and Discussions for "Motivational learning biases are differentially modulated by genetic determinants of striatal and prefrontal dopamine function"

### Supplementary Methods

**Supplementary Table S1. Demographic description of the samples**

| | Cohort 1 | Cohort 2 | Cohort 3 |
|---|---|---|---|
| Gender (N Women/Men) | 43/44 | 48/47 | 55/44 |
| Age in years (M +/- SD) | 24.6 +/- 3.1 | 24.6 +/- 2.8 | 25.2 +/- 2.6 |
| Age range in years | 19 - 36 | 20 - 33 | 20 - 34 |

N = number, M = mean, SD = standard deviation.

##### Trial-by-trial behavioral analysis of *go* responses

To account for both between- and within-subjects variability, individual *go/no-go* responses (1 = *go*, 0 = *no-go*), were analyzed with logistic mixed-level models using the lme4 package in R 3.5.0 [1,2]. The model included all main effects and interactions of the factors action (1 = required *go*, 0 = required *no-go*), valence (1 = *win*, 0 = *avoid losing*) and time (0 = first half, 1 = second half) and a full random effects structure [3-5]. A model containing all interactions of the genotypes with the task conditions did not converge. Therefore, we reduced the model according to our ANCOVA results and tested only the effects of each genotype on the time-dependent valence effect on individual *go/no-go* responses (*genotype x valence x time*).

### Supplementary Results

#### Task effects

In our third independent cohort, we replicated previous results of studies using the same task [3,6-17]. For an overview of the statistics, see Supplementary Table S2. We observed the main effects of *action* ($F_{1,98} = 64.55$, $p < .001$), reflecting the overall *action* bias, and of *time* ($F_{1,98} = 242.48$, $p < .001$), reflecting learning from the first to the second half of the experiment. Moreover, we observed the known interaction of *action* x *valence* ($F_{1,98} = 30.32$, $p < .001$) that was driven by the comparatively low performance in the *no-go to win* condition. Furthermore, we observed a main effect of valence ($F_{1,98} = 6.16$, $p = .015$), and interactions of time with valence ($F_{1,98} = 6.52$, $p = .012$) and time with action ($F_{1,98} = 40.23$, $p < .001$).

**Supplementary Table S2.** An overview of the statistics in the third cohort

| effect | df | *F* | *p* |
|---|---|---|---|
| time | 1.98 | 242.84 | **<.001** |
| action | 1.98 | 64.55 | **<.001** |
| valence | 1.98 | 6.16 | **.015** |
| time * action | 1.98 | 46.23 | **<.001** |
| time * valence | 1.98 | 6.52 | **.012** |
| action * valence | 1.98 | 30.32 | **<.001** |
| time * action * valence | 1.98 | 2.80 | .097 |

Time (1st/2nd half), action (go/no-go), and valence (win/avoid losing) as within-subject factors. Boldface type: $p < .05$.

**Supplementary Table S3.** An overview of the ANCOVA results in the whole cohort.

| effect | df | *F* | *p* |
|---|---|---|---|
| time | 1, 271 | 9.43 | **.002** |
| time * cohort2 | 1, 271 | 1.80 | .181 |
| time * cohort3 | 1, 271 | 2.65 | .105 |
| time * gender | 1, 271 | 0.18 | .676 |
| time * age | 1, 271 | 0.60 | .439 |
| time * TAQ | 1, 271 | 11.08 | **.001** |
| time * COMT | 2, 271 | 2.51 | .083 |
| time * TAQ * COMT | 2, 271 | 2.39 | .093 |
| action | 1, 271 | 0.31 | .582 |
| action * cohort2 | 1, 271 | 1.02 | .313 |
| action * cohort3 | 1, 271 | 1.01 | .316 |
| action * gender | 1, 271 | 0.02 | .884 |
| action * age | 1, 271 | 1.06 | .304 |
| action * TAQ | 1, 271 | 0.01 | .937 |
| action * COMT | 2, 271 | 0.62 | .539 |
| action * TAQ * COMT | 2, 271 | 0.14 | .872 |
| valence | 1, 271 | 0.01 | .922 |
| valence * cohort2 | 1, 271 | 0.07 | .793 |
| valence * cohort3 | 1, 271 | 0.27 | .605 |
| valence * gender | 1, 271 | 0.55 | .459 |
| valence * age | 1, 271 | 0.58 | .448 |
| valence * TAQ | 1, 271 | 0.00 | .962 |
| valence * COMT | 2, 271 | 0.47 | .627 |
| valence * TAQ * COMT | 2, 271 | 2.20 | .112 |
| time * action | 1, 271 | 0.06 | .814 |
| time * action * cohort2 | 1, 271 | 1.16 | .282 |
| time * action * cohort3 | 1, 271 | 0.31 | .581 |
| time * action * gender | 1, 271 | 1.67 | .197 |
| time * action * age | 1, 271 | 0.10 | .754 |
| time * action * TAQ | 1, 271 | 11.94 | **.001** |
| time * action * COMT | 2, 271 | 0.22 | .801 |
| time * action * TAQ * COMT | 2, 271 | 0.93 | .398 |
| time * valence | 1, 271 | 0.64 | .425 |
| time * valence * cohort2 | 1, 271 | 0.02 | .877 |
| time * valence * cohort3 | 1, 271 | 1.18 | .279 |
| time * valence * gender | 1, 271 | 0.32 | .573 |
| time * valence * age | 1, 271 | 0.19 | .660 |
| time * valence * TAQ | 1, 271 | 3.07 | .081 |
| time * valence * COMT | 2, 271 | 0.87 | .419 |
| time * valence * TAQ * COMT | 2, 271 | 0.33 | .719 |

Continued on next page

**Supplementary Table S3** (continued)

| effect | df | *F* | *p* |
|---|---|---|---|
| action * valence | 1, 271 | 0.88 | .348 |
| action * valence * cohort2 | 1, 271 | 0.44 | .508 |
| action * valence * cohort3 | 1, 271 | 1.21 | .273 |
| action * valence * gender | 1, 271 | 0.44 | .508 |
| action * valence * age | 1, 271 | 0.29 | .593 |
| action * valence * TAQ | 1, 271 | 0.56 | .457 |
| action * valence * COMT | 2, 271 | 0.11 | .894 |
| action * valence * TAQ * COMT | 2, 271 | 0.48 | .622 |
| time * action * valence | 1, 271 | 2.77 | .097 |
| time * action * valence * cohort2 | 1, 271 | 2.47 | .117 |
| time * action * valence * cohort3 | 1, 271 | 0.67 | .416 |
| time * action * valence * gender | 1, 271 | 0.04 | .843 |
| time * action * valence * age | 1, 271 | 2.79 | .096 |
| time * action * valence * TAQ | 1, 271 | 11.18 | **.001** |
| time * action * valence * COMT | 2, 271 | 2.96 | .053 |
| time * action * valence * TAQ * COMT | 2, 271 | 0.01 | .987 |
| cohort2 | 1, 271 | 6.77 | **.010** |
| cohort3 | 1, 271 | 2.60 | .108 |
| gender | 1, 271 | 3.55 | .061 |
| age | 1, 271 | 6.67 | **.010** |
| TAQ | 1, 271 | 0.97 | .327 |
| COMT | 2, 271 | 2.53 | .082 |
| TAQ * COMT | 2, 271 | 2.03 | .133 |

Time (1st/2nd half), action (go/no-go), and valence (win/avoid losing) as within-subject factors, genotypes as between-subject factors, cohorts (three cohorts represented in two dichotomous dummy-coded variables for cohort 2 and 3), age and gender as covariates. Boldface type: $p < .05$.

**Supplementary Table S4.** An overview of the *post hoc* comparisons in the whole cohort

| variable | TaqIA | N | Mean | SD | *t* | df | *p* | Cohen's *d* |
|---|---|---|---|---|---|---|---|---|
| all conditions 1st half | A1 carriers | 99 | 76.18 | 14.95 | -0.15 | 173.28 | 0.879 | |
| | A2 homozygotes | 182 | 76.45 | 12.51 | | | | |
| all conditions 2nd half | A1 carriers | 99 | 86.55 | 16.95 | -2.21 | 158.90 | **0.028** | 0.35 |
| | A2 homozygotes | 182 | 90.85 | 12.69 | | | | |
| go conditions 1st half | A1 carriers | 99 | 84.28 | 14.30 | -1.18 | 146.28 | 0.241 | |
| | A2 homozygotes | 182 | 86.16 | 9.52 | | | | |
| go conditions 2nd half | A1 carriers | 99 | 91.75 | 12.72 | -1.51 | 169.00 | 0.133 | |
| | A2 homozygotes | 182 | 94.00 | 10.32 | | | | |
| nogo conditions 1st half | A1 carriers | 99 | 68.08 | 20.57 | 0.55 | 279.00 | 0.583 | |
| | A2 homozygotes | 182 | 66.73 | 19.18 | | | | |
| nogo conditions 2nd half | A1 carriers | 99 | 81.35 | 24.06 | -2.28 | 164.61 | **0.024** | 0.29 |
| | A2 homozygotes | 182 | 87.70 | 18.88 | | | | |
| go to win condition 1st half | A1 carriers | 99 | 87.78 | 21.18 | -1.68 | 147.19 | 0.094 | |
| | A2 homozygotes | 182 | 91.78 | 14.23 | | | | |
| go to avoid losing condition 1st half | A1 carriers | 99 | 80.77 | 14.13 | 0.14 | 163.80 | 0.891 | |
| | A2 homozygotes | 182 | 80.55 | 11.02 | | | | |
| no-go to win condition 1st half | A1 carriers | 99 | 62.39 | 31.99 | 0.96 | 279.00 | 0.34 | |
| | A2 homozygotes | 182 | 58.57 | 31.99 | | | | |
| no-go to avoid losing condition 1st half | A1 carriers | 99 | 73.77 | 16.48 | -0.58 | 279.00 | 0.564 | |
| | A2 homozygotes | 182 | 74.89 | 15.00 | | | | |
| go to win condition 2nd half | A1 carriers | 99 | 93.67 | 17.63 | -1.02 | 279.00 | 0.309 | |
| | A2 homozygotes | 182 | 95.60 | 13.70 | | | | |
| go to avoid losing condition 2nd half | A1 carriers | 99 | 89.83 | 14.93 | -1.49 | 162.35 | 0.139 | |
| | A2 homozygotes | 182 | 92.40 | 11.51 | | | | |
| no-go to win condition 2nd half | A1 carriers | 99 | 72.56 | 38.38 | -2.06 | 175.98 | **0.041** | 0.26 |
| | A2 homozygotes | 182 | 81.94 | 32.74 | | | | |
| no-go to avoid losing condition 2nd half | A1 carriers | 99 | 90.13 | 16.16 | -1.76 | 168.82 | 0.081 | |
| | A2 homozygotes | 182 | 93.46 | 13.09 | | | | |
| slope all conditions | A1 carriers | 99 | 10.37 | 7.79 | -3.72 | 279.00 | **<0.001** | 0.47 |
| | A2 homozygotes | 182 | 14.40 | 9.14 | | | | |
| slope go conditions | A1 carriers | 99 | 7.47 | 9.83 | -0.32 | 279.00 | 0.748 | |
| | A2 homozygotes | 182 | 7.84 | 8.59 | | | | |
| slope nogo conditions | A1 carriers | 99 | 13.27 | 12.81 | -4.56 | 279.00 | **<0.001** | 0.58 |
| | A2 homozygotes | 182 | 20.97 | 13.88 | | | | |
| slope go to win condition | A1 carriers | 99 | 5.89 | 13.68 | 1.31 | 158.91 | 0.191 | |
| | A2 homozygotes | 182 | 3.83 | 10.25 | | | | |
| slope go to avoid losing condition | A1 carriers | 99 | 9.06 | 15.08 | -1.72 | 279.00 | 0.087 | |
| | A2 homozygotes | 182 | 11.85 | 11.77 | | | | |
| slope no-go to win condition | A1 carriers | 99 | 10.17 | 22.94 | -4.41 | 221.84 | **<0.001** | 0.54 |
| | A2 homozygotes | 182 | 23.37 | 25.74 | | | | |
| slope no-go to avoid losing condition | A1 carriers | 99 | 16.36 | 12.14 | -1.42 | 279.00 | 0.158 | |
| | A2 homozygotes | 182 | 18.57 | 12.65 | | | | |

Boldface type: $p < .05$.

#### Trial-by-trial behavioral analysis of *go* responses

A summary of the trial-by-trial behavioral analysis is presented in Figure S1. Subjects successfully learned the task and were able to adjust *go* responses to the required action as evidenced by a significantly positive effect of *action* ($z$ = 31.16, $p$ < .001). A significantly negative effect of *time* ($z$ = -18.58, $p$ < .001) indicated that they started the experiment with a *go* bias, but could improve over time as evident in a significant *action* x *time* interaction ($z$ = 21.03, $p$ < .001). As expected, there was a motivational bias in *go* responding revealed by a significant *valence* effect ($z$ = 5.44, $p$ < .001) akin to the *action* x *valence* interaction for accuracy and hence our main effect of interest. This *valence* effect was stronger for *go* cues as evident in a significantly positive *action* x *valence* interaction ($z$ = 6.85, $p$ < .001).

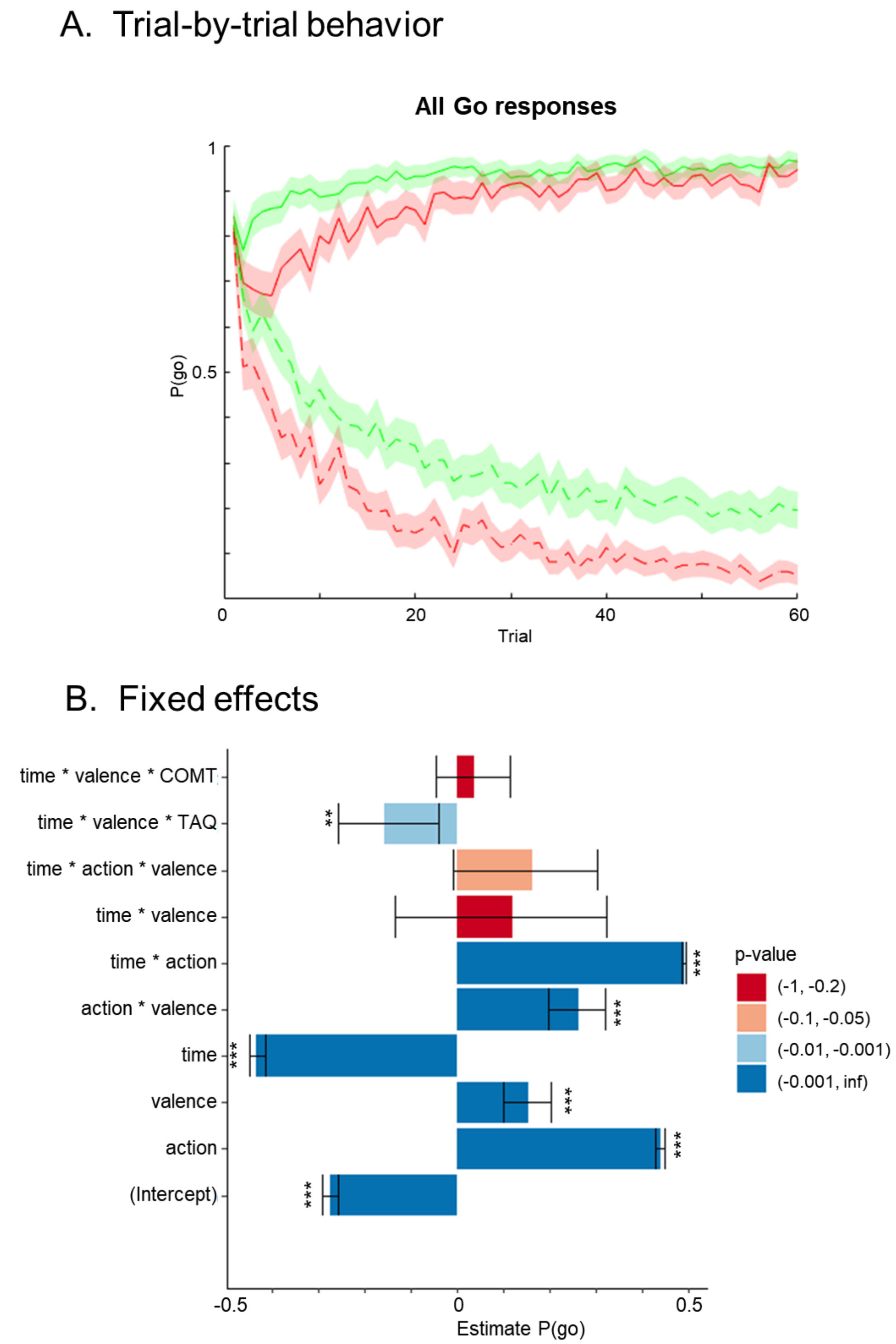


**Fig. S1. Trial-by-trial behavior.** *(A)* Trial-by-trial proportions of *go* responses (±SEM) to *go* cues (solid lines) and *no-go* cues (dashed lines) across cue types. Participants made more *go* responses to *win* vs. *avoid losing* cues (i.e. green lines are above red lines), reflecting the motivational bias. Overall, they successfully learned whether to make a *go* response or not (proportion of *go* responses increases for *go* cues and decreases for *no-go* cues). *(B)* Logistic mixed model estimates of the probability of *go* responses. Fixed effect estimates and 95% confidence interval (CI) are plotted on probability scale. *$p < 0.05$, **$p < 0.01$, ***$p < 0.001$. Adapted scripts of Swart et al. [3] were used to generate figures.

In line with the ANCOVA results on accuracy, DRD2 TaqIA A1 carriers showed a more pronounced effect of cue valence on *go* responding specifically at the second half of the experiment (Figure 1C), which was reflected by a significantly negative *genotype* x *valence* x *time* interaction ($z = 2.62$, $p = .009$). For the COMT SNP no interaction with the time-dependent valence effect on individual *go/no-go* responses could be observed ($p = .381$; see Figure S2).

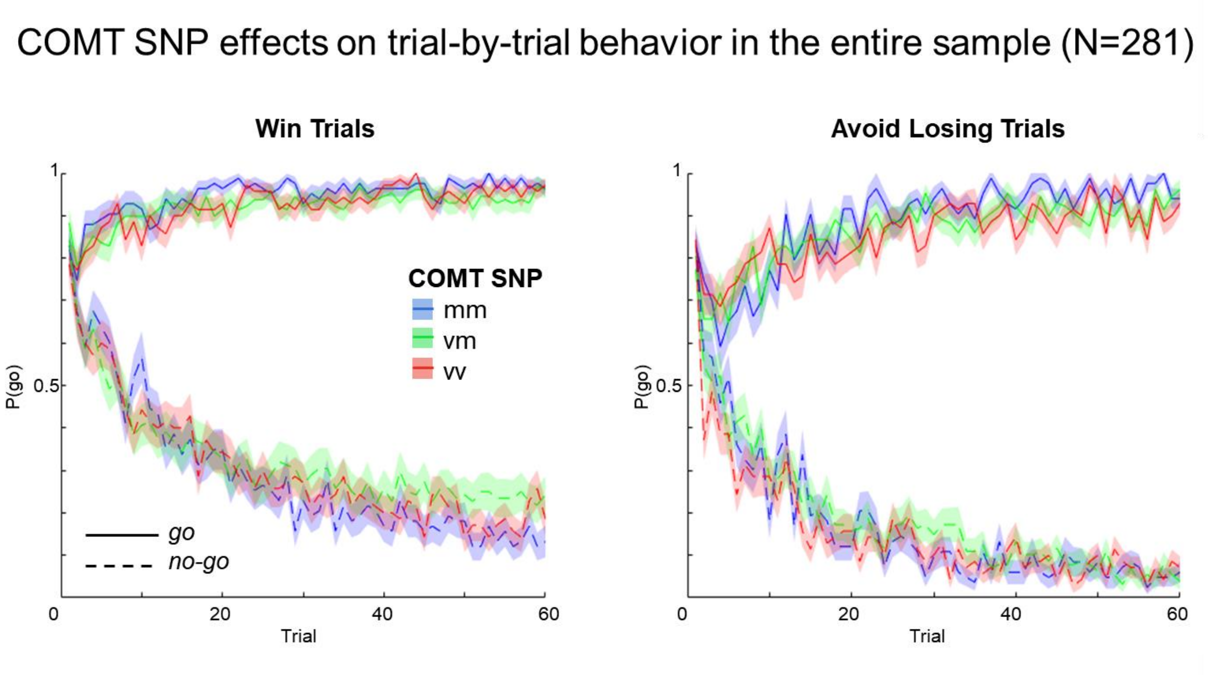


**Fig. S2. Effects of COMT genotype on *go* responses in the entire sample.** Trial-by-trial proportions of go responses (±SEM) to go cues (solid lines) and no-go cues (dashed lines) across cue types. Win and avoid losing condition seperately and colors depict COMT genotypes. Adapted scripts of Swart et al. [3] were used to generate figures.

##### Analysis of reaction times

As in our previous study [17], the analysis of the reaction times (RTs) in the *go* conditions revealed a *valence* effect ($F_{1,275} = 32.28$, $p < .001$) with faster RTs in the *win* (M +/- SD = 542 +/- 99 ms) compared to the *avoid losing* (M +/- SD = 563 +/- 107 ms) condition (paired *t*-test: $t_{280} = -5.76$, $p < .001$). No effects of *time* or *genotype* were observed ($p > .221$).

##### Analysis of false responses

As a control, we investigated the number of trials where participants responded incorrectly (i.e., left when the target was on the right side of the display or vice versa). False response rates were very low (M +/- SD = 0.5+/-2.0%) and did not differ between genotype groups (TaqIA: Mann-Whitney *U*-test $p = .403$; COMT: Kruskal-Wallis *H*-test $p = .187$), thus ruling out the possibility that genotype effects could be driven by differences in the target detection performance.

### Supplementary Discussion

##### Potential molecular mechanisms of the DRD2/ANKK1 TaqIA polymorphism

ANKK1, also known as receptor interacting protein 5 [18], codes a potential kinase the function of which is largely unknown. Ankyrin repeat domains are frequent sequential motives that regulate protein-protein interactions and influence the stability, folding and unfolding, as well as the binding properties of proteins [19]. As the *DRD2* and *ANKK1* genes are closely linked [20,21], it has been suggested that genetic variations in linkage disequilibrium (LD) with TaqIA might explain the observed relationship between the SNP and alterations of human dopaminergic neurotransmission. Indeed the *DRD2/ANKK1* TaqIA polymorphism is in LD with several polymorphisms on the DRD2 gene of which especially the C957T polymorphism has received considerable attention [22-26]. Until now the number of studies investigating *ANKK1* and *DRD2* polymorphisms together is small and their results are complex. While some postulate the highest risk by the combination of both variants of TaqIA and C957T polymorphism [22,27-32], others show that TaqIA effects are carried by the C957T polymorphism [33], and yet others suggest a modulatory effect of the *ANKK1* gene independent of the DRD2 gene [34-37]. The latter, for example, was confirmed in studies that investigated different variants in the *NCAM1-TTC12-ANKK1-DRD2* region and found the highest associations between addiction disorder and polymorphisms on the *ANKK1* gene [34,35]. But also PET studies showing effects of TaqIA but not C957T polymorphism on DA activity affirm this assumption [36,37]. However, one should be careful with associations of *ANKK1* variants, DRD2 expression and a phenotype [38]. Anyhow current studies point out that *ANKK1* might be involved in a regulatory signaling cascade, that influences the dopaminergic system and especially D2 receptors, what could potentially be modulated by the TaqIA polymorphism [18,39,40] (for a review, see [21]).

##### COMT-dependent clearance of cortical dopamine

The role of COMT in dopamine clearance has been subject to extensive research since the first studies suggesting a role for the COMT Val108/158Met polymorphism in PFC function [41,42]. Converging evidence from animal studies and human *post mortem* investigations suggests that COMT is primarily important for dopamine inactivation in brain regions where the dopamine transporter (DAT) is sparsely expressed, especially the prefrontal cortex (PFC) [43-45].

Despite the wide agreement regarding the preferential role for COMT in prefrontal versus striatal dopamine inactivation, the contributing cellular mechanisms have been subject to considerable debate. Particularly, the question whether COMT can be active in the extracellular space or whether a form of DAT-independent dopamine uptake is required before COMT-dependent dopamine inactivation, has yielded conflicting results in previous research. Most of the debate concerns membrane-bound COMT (MB-COMT), the predominant form in neurons and glial cells [46]. In a frequently cited publication, Chen et al. [47] reported a series of experiments from which they deduced that neuronal MB-COMT is oriented extracellularly and exhibits extracellular enzymatic activity. The authors could demonstrate the expression of COMT in neurons, but stainings were performed in permeabilized cells and did thus not provide information about intra- versus extracellular localization of epitopes. On the other hand, live staining experiments have shown that COMT epitopes can only be detected after permeabilization of the cell membrane [48,49].

Even if one was to assume the occasional presence of extracellular COMT catalytic domains, an important question is whether they could be enzymatically active. COMT activity depends on magnesium ($Mg^{2+}$) ions, and it is inhibited by calcium ($Ca^{2+}$) [50]. Intracellular $Mg^{2+}$ reaches a concentration of 5 to 20 mmol/l, whereas extracellular $Mg^{2+}$ concentrations range from 0.6 to 1.0 mmol/l [51]. Conversely, $Ca^{2+}$ is the predominant divalent cation in the extracellular space (2.2 – 2.6 mmol/l), while its intracellular concentration is low. In their article, Chen et al. [47] described that presumably extracellular COMT activity was measured in a medium containing Tris buffer and magnesium chloride, which is poorly comparable with the natural extracellular milieu where the catalytic domain of COMT would most likely be inhibited by the two-to-fourfold higher concentrations of $Ca^{2+}$ compared to $Mg^{2+}$. In summary, evidence for physiologically relevant extracellular COMT activity must be considered questionable. Therefore, a DAT-independent uptake mechanism must contribute to COMT-dependent dopamine clearance in the PFC. One study suggests that norepinephrine transporter (NET) likely contributes to this uptake [43], but additional candidate transport proteins may also play a role in DAT-independent dopamine uptake (for a discussion, see [48]).

18 Garrido E, Palomo T, Ponce G, Garcia-Consuegra I, Jimenez-Arriero MA, Hoenicka J. The ANKK1 protein associated with addictions has nuclear and cytoplasmic

localization and shows a differential response of Ala239Thr to apomorphine. Neurotox Res. 2011;20(1):32-9.
19 Li J, Mahajan A, Tsai MD. Ankyrin repeat: a unique motif mediating protein-protein interactions. Biochemistry. 2006;45(51):15168-78.
20 Neville MJ, Johnstone EC, Walton RT. Identification and characterization of ANKK1: a novel kinase gene closely linked to DRD2 on chromosome band 11q23.1. Hum Mutat. 2004;23(6):540-5.
21 Ponce G, Perez-Gonzalez R, Aragues M, Palomo T, Rodriguez-Jimenez R, Jimenez-Arriero MA, et al. The ANKK1 kinase gene and psychiatric disorders. Neurotox Res. 2009;16(1):50-9.
22 Doehring A, Hentig N, Graff J, Salamat S, Schmidt M, Geisslinger G, et al. Genetic variants altering dopamine D2 receptor expression or function modulate the risk of opiate addiction and the dosage requirements of methadone substitution. Pharmacogenet Genomics. 2009;19(6):407-14.
23 Ritchie T, Noble EP. Association of seven polymorphisms of the D2 dopamine receptor gene with brain receptor-binding characteristics. Neurochemical research. 2003;28(1):73-82.
24 Richter A, Barman A, Wustenberg T, Soch J, Schanze D, Deibele A, et al. Behavioral and Neural Manifestations of Reward Memory in Carriers of Low-Expressing versus High-Expressing Genetic Variants of the Dopamine D2 Receptor. Front Psychol. 2017;8:654.
25 Fossella J, Green AE, Fan J. Evaluation of a structural polymorphism in the ankyrin repeat and kinase domain containing 1 (ANKK1) gene and the activation of executive attention networks. Cognitive, affective & behavioral neuroscience. 2006;6(1):71-8.
26 Duan J, Wainwright MS, Comeron JM, Saitou N, Sanders AR, Gelernter J, et al. Synonymous mutations in the human dopamine receptor D2 (DRD2) affect mRNA stability and synthesis of the receptor. Hum Mol Genet. 2003;12(3):205-16.
27 Hirvonen MM, Laakso A, Nagren K, Rinne JO, Pohjalainen T, Hietala J. C957T polymorphism of dopamine D2 receptor gene affects striatal DRD2 in vivo availability by changing the receptor affinity. Synapse. 2009;63(10):907-12.
28 Morton LM, Wang SS, Bergen AW, Chatterjee N, Kvale P, Welch R, et al. DRD2 genetic variation in relation to smoking and obesity in the Prostate, Lung, Colorectal, and Ovarian Cancer Screening Trial. Pharmacogenet Genomics. 2006;16(12):901-10.
29 Ponce G, Hoenicka J, Jimenez-Arriero MA, Rodriguez-Jimenez R, Aragues M, Martin-Sune N, et al. DRD2 and ANKK1 genotype in alcohol-dependent patients with psychopathic traits: association and interaction study. Br J Psychiatry. 2008;193(2):121-5.
30 Swagell CD, Lawford BR, Hughes IP, Voisey J, Feeney GF, van Daal A, et al. DRD2 C957T and TaqIA genotyping reveals gender effects and unique low-risk and high-risk genotypes in alcohol dependence. Alcohol Alcohol. 2012;47(4):397-403.
31 Voisey J, Swagell CD, Hughes IP, van Daal A, Noble EP, Lawford BR, et al. A DRD2 and ANKK1 haplotype is associated with nicotine dependence. Psychiatry Res. 2012;196(2-3):285-9.
32 Yang BZ, Kranzler HR, Zhao H, Gruen JR, Luo X, Gelernter J. Haplotypic variants in DRD2, ANKK1, TTC12, and NCAM1 are associated with comorbid alcohol and drug dependence. Alcoholism, clinical and experimental research. 2008;32(12):2117-27.
33 Frank MJ, Hutchison K. Genetic contributions to avoidance-based decisions: striatal D2 receptor polymorphisms. Neuroscience. 2009;164(1):131-40.
34 Dick DM, Wang JC, Plunkett J, Aliev F, Hinrichs A, Bertelsen S, et al. Family-based association analyses of alcohol dependence phenotypes across DRD2 and neighboring gene ANKK1. Alcoholism, clinical and experimental research. 2007;31(10):1645-53.

35 Gelernter J, Yu Y, Weiss R, Brady K, Panhuysen C, Yang BZ, et al. Haplotype spanning TTC12 and ANKK1, flanked by the DRD2 and NCAM1 loci, is strongly associated to nicotine dependence in two distinct American populations. Hum Mol Genet. 2006;15(24):3498-507.

36 Laakso A, Pohjalainen T, Bergman J, Kajander J, Haaparanta M, Solin O, et al. The A1 allele of the human D2 dopamine receptor gene is associated with increased activity of striatal L-amino acid decarboxylase in healthy subjects. Pharmacogenet Genomics. 2005;15(6):387-91.

37 Joutsa J, Hirvonen MM, Arponen E, Hietala J, Kaasinen V. DRD2-related TaqIA genotype is associated with dopamine release during a gambling task. J Addict Med. 2014;8(4):294-5.

38 Lucht M, Rosskopf D. Comment on "Genetically determined differences in learning from errors". Science. 2008;321(5886):200; author reply 00.

39 Hoenicka J, Quinones-Lombrana A, Espana-Serrano L, Alvira-Botero X, Kremer L, Perez-Gonzalez R, et al. The ANKK1 gene associated with addictions is expressed in astroglial cells and upregulated by apomorphine. Biological psychiatry. 2010;67(1):3-11.

40 Ponce G, Quinones-Lombrana A, Martin-Palanco NG, Rubio-Solsona E, Jimenez-Arriero MA, Palomo T, et al. The Addiction-Related Gene Ankk1 is Oppositely Regulated by D1R- and D2R-Like Dopamine Receptors. Neurotox Res. 2016;29(3):345-50.

41 Weinberger DR, Egan MF, Bertolino A, Callicott JH, Mattay VS, Lipska BK, et al. Prefrontal neurons and the genetics of schizophrenia. Biological psychiatry. 2001;50(11):825-44.

42 Egan MF, Goldberg TE, Kolachana BS, Callicott JH, Mazzanti CM, Straub RE, et al. Effect of COMT Val108/158 Met genotype on frontal lobe function and risk for schizophrenia. Proceedings of the National Academy of Sciences of the United States of America. 2001;98(12):6917-22.

43 Kaenmaki M, Tammimaki A, Myohanen T, Pakarinen K, Amberg C, Karayiorgou M, et al. Quantitative role of COMT in dopamine clearance in the prefrontal cortex of freely moving mice. J Neurochem. 2010;114(6):1745-55.

44 Yavich L, Forsberg MM, Karayiorgou M, Gogos JA, Mannisto PT. Site-specific role of catechol-O-methyltransferase in dopamine overflow within prefrontal cortex and dorsal striatum. The Journal of neuroscience : the official journal of the Society for Neuroscience. 2007;27(38):10196-209.

45 Matsumoto M, Weickert CS, Akil M, Lipska BK, Hyde TM, Herman MM, et al. Catechol O-methyltransferase mRNA expression in human and rat brain: evidence for a role in cortical neuronal function. Neuroscience. 2003;116(1):127-37.

46 Myohanen TT, Schendzielorz N, Mannisto PT. Distribution of catechol-O-methyltransferase (COMT) proteins and enzymatic activities in wild-type and soluble COMT deficient mice. J Neurochem. 2010;113(6):1632-43.

47 Chen J, Song J, Yuan P, Tian Q, Ji Y, Ren-Patterson R, et al. Orientation and cellular distribution of membrane-bound catechol-O-methyltransferase in cortical neurons: implications for drug development. J Biol Chem. 2011;286(40):34752-60.

48 Schott BH, Frischknecht R, Debska-Vielhaber G, John N, Behnisch G, Duzel E, et al. Membrane-Bound Catechol-O-Methyl Transferase in Cortical Neurons and Glial Cells is Intracellularly Oriented. Front Psychiatry. 2010;1:142.

49 Ulmanen I, Peranen J, Tenhunen J, Tilgmann C, Karhunen T, Panula P, et al. Expression and intracellular localization of catechol O-methyltransferase in transfected mammalian cells. Eur J Biochem. 1997;243(1-2):452-9.

50 Sparta M, Alexandrova AN. How metal substitution affects the enzymatic activity of catechol-o-methyltransferase. PLoS One. 2012;7(10):e47172.
51 Jahnen-Dechent W, Ketteler M. Magnesium basics. Clin Kidney J. 2012;5(Suppl 1):i3-i14.
